# Airway fibroblasts drive WNT5B-dependent mast cell progenitor proliferation

**DOI:** 10.64898/2026.11.16.766533

**Authors:** Tahereh Derakhshan, Kenshiro Matsuda, Eleanor Hollers, Rizza U. Santos, Camden A. New, Suppawat Kongthong, Radomir Kratchmarov, Alexander Perniss, Regan W. Bergmark, Stella E. Lee, Alice Z. Maxfield, Rachel E. Roditi, Kathleen M. Buchheit, Tanya M. Laidlaw, Joshua A. Boyce, Daniel F. Dwyer

**Affiliations:** Jeff and Penny Vinik Center for Allergic Disease Research, Division of Allergy and Clinical Immunology, Mass General Brigham, Boston, Massachusetts, U.S.A.; Harvard Medical School, Boston, Massachusetts, U.S.A.; Division of Rheumatology and Immunology, Mass General Brigham, Boston, Massachusetts, U.S.A.; Department of Otolaryngology – Head and Neck Surgery, Mass General Brigham, Boston, Massachusetts, U.S.A.

## Abstract

Mast cells (MCs) are tissue-resident granulocytes that play key roles in human type 2 (T2) inflammatory disease through release of preformed and newly generated pro-inflammatory mediators^1^. Paradoxically, while MCs are abundantly present during T2 inflammation^2–5^, MC progenitors (MCps) are among the rarest circulating cells and proliferate slowly^6^. Thus, it is unclear how such scarce progenitors can sustain the marked MC hyperplasia observed in T2 disease. To address this, we conducted single-cell multiomics profiling on paired peripheral blood progenitors and nasal polyp MCs from patients undergoing polypectomy. This approach defined cell surface and intracellular markers linking circulating MCps to a recently identified proliferative MC subset^5^. In vitro co-culture studies revealed that sinonasal fibroblasts markedly enhanced both MCp proliferation and maturation and identified WNT5B as a novel fibroblast-derived mitogen responsible for amplifying MCp proliferation. Together, these findings establish fibroblasts as a key regulator of MC hyperplasia during tissue inflammation.

## Main Text

To assess the relationship between circulating MCps and mature MCs within nasal polyps, CD45^+^ Lineage (CD3, CD11b, CD11c, CD14, CD19, CD56, CCR3)-negative cells were sort-purified from polyp tissue obtained from eight donors undergoing sinonasal surgery to treat chronic rhinosinusitis (CRS) with nasal polyps (CRSwNP), in parallel with autologous CD34^+^ peripheral blood mononuclear cells (PBMCs) collected at the time of surgery (Fig. 1A, Fig. S1A-B). The donors included a mix of aspirin-tolerant CRSwNP patients and patients with aspirin-exacerbated respiratory disease (AERD), a severe endotype of CRSwNP associated with rapid polyp regrowth, co-morbid asthma, and increased MC burden within polyp tissue^7^. Cells were stained with hashtag antibodies to designate tissue of origin and oligo-tagged antibodies for a select panel of cell surface markers (CD117, Integrin αE, Integrin β7, FcεR1α, CD38, CD49d, CD133, CD131, CXCR4, CD71, four donors) alone or in conjunction with a cocktail of 134 antibodies (four donors) prior to pooling for single-cell RNA-seq (scRNA-seq). Cells expressing either *CD34* or MC protease-encoding genes (*TPSAB1, CPA3*) were computationally isolated for downstream analysis, excluding any clusters co-expressing *TPSAB1* and lymphocyte lineage-associated transcripts (*MZB1*, *CD3E*) (Fig. S1C-D).

**Figure 1:**
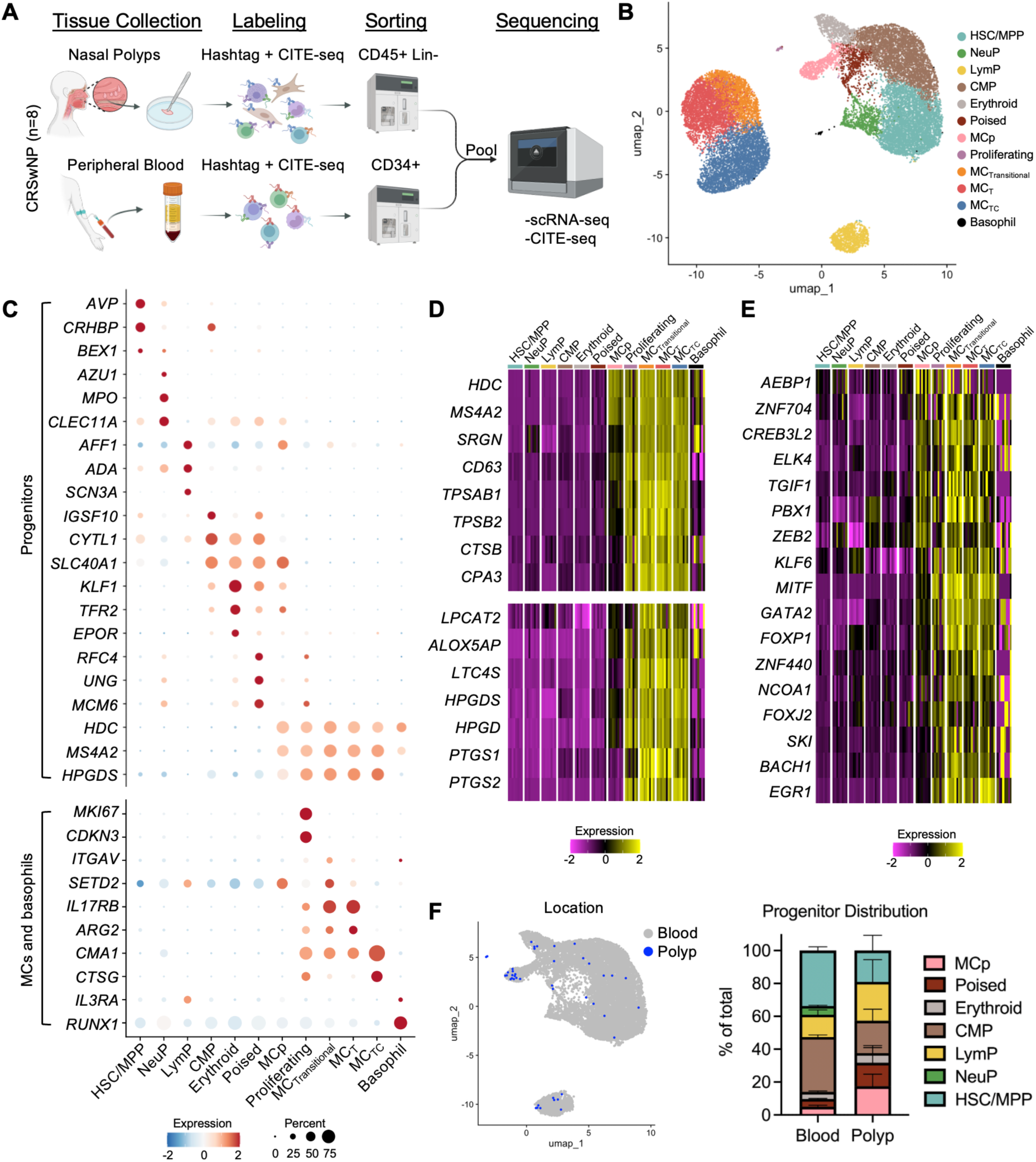
Single-cell characterization of circulating progenitor populations. A) Schematic representation for isolating lineage negative cells from nasal polyp tissue and autologous circulating progenitor cells for multiomics analysis. B) Uniform manifold approximation and projection (UMAP) embedding of circulating progenitor and mature tissue-resident populations. C) Marker gene expression profiles for circulating progenitors (top) and mature MCs and basophils (bottom). D) Donor-averaged row-normalized heatmap showing expression of MC activation- and granule-associated transcripts (top) and eicosanoid biosynthetic enzyme-encoding transcripts (bottom) enriched in circulating MCp relative to other progenitor populations. E) Donor-averaged row-normalized heatmap showing expression of transcription factor-encoding transcripts enriched in circulating MCp relative to other progenitor populations. F) UMAP display (left) and quantification (right) of circulating progenitor populations identified via hashtag as originating in peripheral blood (grey) or polyp (blue). Scale bars indicate z score. For all heatmaps, Padj < 0.05 and Log2FoldChange > 0.5 for MCp relative to circulating progenitors. Columns indicate cluster-averaged expression for each individual donor.

The remaining cells were re-clustered, after which differential gene expression analysis identified canonical population-associated transcripts enriched in each cluster, which were then used for cluster annotation based on prior studies of human progenitor cells^8,9^ (Fig. 1B-C, Fig. S2A, Table S1). Clusters were consistently observed across donors, with tissue mast cell and progenitor populations readily distinguished based on expression of *TPSAB1* or *CD34* and by tissue of origin (Fig. S2B-D). Within the blood-associated clusters, we identified seven circulation-enriched progenitor clusters, including hematopoietic stem cells / multipotent progenitors (HSC/MPP), neutrophil progenitors (NeuP), lymphoid progenitors (LymP), common myeloid progenitors (CMP), erythroid-lineage progenitors (Erythroid), and MCps. Two clusters were enriched for cell cycle-associated genes and, following cell cycle analysis, were designated as poised (enriched for S but not G2M signatures) and proliferating (enriched for both S and G2M) (Fig. S2E). Tissue MC subsets were distinguished as previously reported based on expression of transcripts associated with subepithelial MC_TC_s (*CMA1*, *CTSG*), intraepithelial MC_T_s (*IL17RB*, *GPR183*), transitional MCs (*ITGAV*, *SETD2*)^5,10^, while basophils were identified based on co-expression of *IL3RA* and *RUNX1* (Fig. 1C, S2A, Table S2).

Consistent with prior studies, MCps were enriched for transcripts associated with MC proteases (*TPSB2*, *CPA3*), granule structure (*SRGN*, *CD63*), and eicosanoid biosynthesis (*LPCAT2*, *ALOX5AP*, *LTC4S*, *HPGDS*) relative to other circulating progenitors, albeit at levels much lower than those observed in mature MC populations (Fig. 1C-D). MCps expressed *HDC* (encoding histidine decarboxylase) and *MS4A2* (encoding the FcεR1 beta chain) at levels similar to their tissue counterparts (Fig. 1D). MCps were further enriched for a large subset of transcription factors (TFs) shared with mature MCs, including both TFs known to be required for MC differentiation (*MITF*, *GATA2*) and other less well-defined TFs shared with tissue MCs (*CREB3L2*, *PBX1*, *ZEB2*)^11,12^, suggesting they may also be important for commitment to the MC lineage (Fig. 1E). Interestingly, while MCps represented less than 5% of the circulating CD34⁺ pool, they were the largest population observed within the polyp, where they comprised nearly 20% of tissue-associated progenitors (Fig. 1F, S2D). Thus, a cell cluster with a transcriptional phenotype consistent with circulating MCps was detected in both the circulation and polyp tissue.

We next analyzed cell surface protein expression to further characterize surface markers that distinguish circulating MCps, identifying a series of candidates significantly enriched in MCps among progenitor subsets (Fig. 2A, Table S3). Of these, many were robustly expressed on mature MC populations, while others appeared more strongly associated with progenitors (CD71, CD131, Integrin β7). We additionally identified candidate negative selection markers that were elevated in other progenitors but were low or absent on MCps (CD45RA, GPR56, HLA-DR-DP-DQ) (Fig. 2B). Prior studies have established that cord blood MCps express high levels of *TFRC* (encoding CD71), and differential expression of CD131 distinguishes bone marrow MCps from myeloid/neutrophil progenitors^13,14^. Flow cytometry analysis of CD34^+^ cells confirmed that FcεR1α^+^CD117^+^ MCps could only be observed within the CD71^+^/CD131^+^ gate (Fig. 2C). We further observed that MCps were enriched for cell-surface expression of CD38, CD84, CLEC12A and CD33 via flow cytometry, while exhibiting reduced expression of CD45RA, GPR56 and MHC-II relative to other circulating progenitors (Fig. S3A-B). Thus, CD71 and CD131 co-expression was used as an additional gating criterion to more cleanly identify MCps for downstream characterization and isolation. CD131 pairs with CD123 (IL-3Rα), CD125 (IL-5Rα), or CD116 (GM-CSFRα) to enable recognition of IL-3, IL-5 or GM-CSF, respectively. As previously reported^15^, MCps expressed IL-3Rα and IL-5Rα, but lacked GM-CSFRα expression (Fig. S3C). MCps were further confirmed to exhibit cell surface expression of CD203c and MS4A3, encoded by *ENPP3* and *MS4A3*, respectively (Fig. S3D). The MCp cluster was significantly enriched for transcripts associated with eosinophils and basophils (*PRG2*, *EPX*, *CLC*) and the megakaryocyte/erythroid lineage (*GATA1*, *KLF1*, *ITGA2B*) (Fig. 2D), likely related to their reported shared lineage development^13^. Apart from *PRG2*, expression of basophil, eosinophil and erythroid lineage transcripts was not observed in tissue-resident MCs, suggesting these transcripts were lost during differentiation (Fig. 2D). Consistent with their transcriptional profile, intracellular staining revealed that MCps contained detectable levels of Galectin-10 (CLC), the protein encoded by *CLC*, and 2D7, a reportedly basophil-specific protein of unknown identity^16^, with significant enrichment for both relative to other circulating CD34^+^ cells (Fig. S3E). No intracellular CLC or 2D7 was detected in cells lacking FcεR1α, indicating these proteins were specific to MCps rather than broadly expressed on progenitor cells (Fig. S3F). MCp expression of both proteins was significantly lower than that observed in circulating basophils (Fig. 2E-G). However, CD71 was absent from basophils (Fig. 2H), confirming that 2D7 and CLC detection in MCps was not the result of basophil contamination. Circulating basophils exhibited bimodal expression of CLC but not 2D7 (Fig. S3G), consistent with recent reports of basophil transcriptional heterogeneity stratified in part by *CLC* expression^17^.

**Figure 2:**
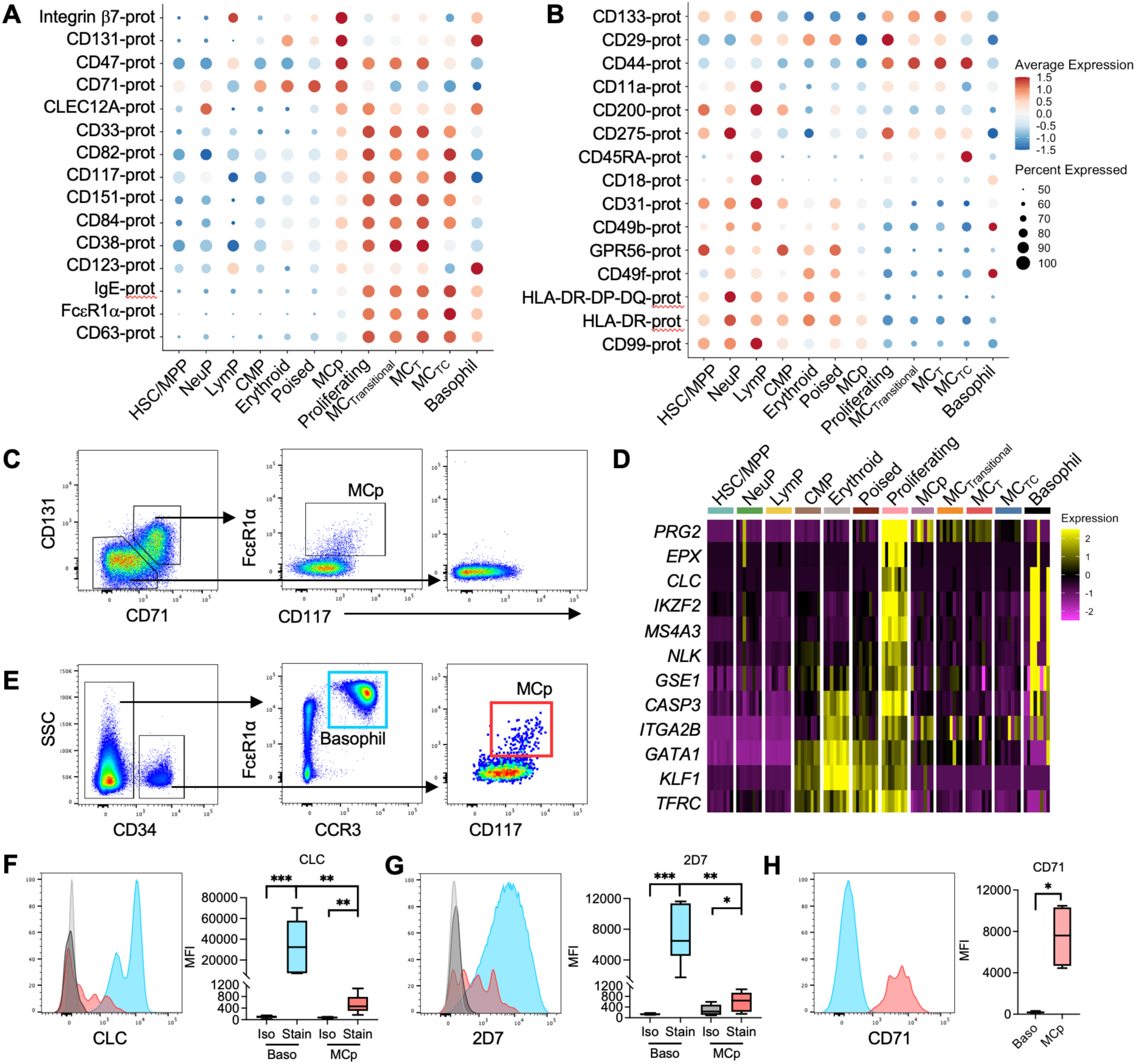
Phenotypic characterization of circulating MCps. A-B) Dot plot showing expression of select cell surface proteins identified via cellular indexing of transcriptomes and epitopes by sequencing (CITE-seq) as enriched (A) or reduced (B) in MCp relative to other CD34^+^ progenitor cells (n=4 donors). C) Flow gating scheme for identifying MCp within the CD131^+^/CD71^+^ population of circulating CD34^+^ cells. D) Transcript expression of select basophil/eosinophil- or erythroid-associated transcripts enriched in MCp relative to other circulating progenitors. E) Gating scheme for identifying MCp versus basophils. F-G) Intracellular staining for CLC (F) and 2D7 (G) in MCp (red) vs circulating basophils (blue) and isotype control staining (light grey, basophil; dark grey, MCp) with quantification (n=6 donors). H) Representative histogram showing CD71 expression in MCp (red) relative to circulating basophils (blue) with quantification (n=4 donors). *, p < 0.05; **, p < 0.01; ***, p < 0.001 (ratio-paired t test for pairwise comparisons, ANOVA for multi-group comparisons). For CITE-seq and transcript data, Padj < 0.05 and Log2FoldChange > 0.5 for MCp relative to other progenitor populations. For dot plots, color denotes cluster-averaged relative expression, size indicates percent of cells with marker detected. For heatmap, columns indicate cluster-averaged expression for each individual donor.

Although cell hashing indicated the presence of MCps within nasal polyp tissue (Fig. 1F), these cells were quite rare, suggesting that the MCp transcriptome is only transiently expressed following recruitment to inflamed tissue. As protein expression lags transcriptional changes, we assessed whether any populations within the nasal polyp MC pool exhibited an MCp-like cell surface phenotype by conducting differential protein analysis of polyp-resident MC subsets. Surprisingly, we observed that proliferating MCs exhibited a distinct cell surface profile relative to other polyp MC populations (Fig. 3A, Table S3). Among these proliferation-associated markers were CD71, previously identified as MCp-associated in the circulation, and CD49d, CD31, CD43 and CD93, all robustly expressed by circulating MCps (Fig. 3B-C). Thus, CITE-seq analysis suggested a potential link between circulating MCps and proliferating MCs within the polyp environment.

**Figure 3:**
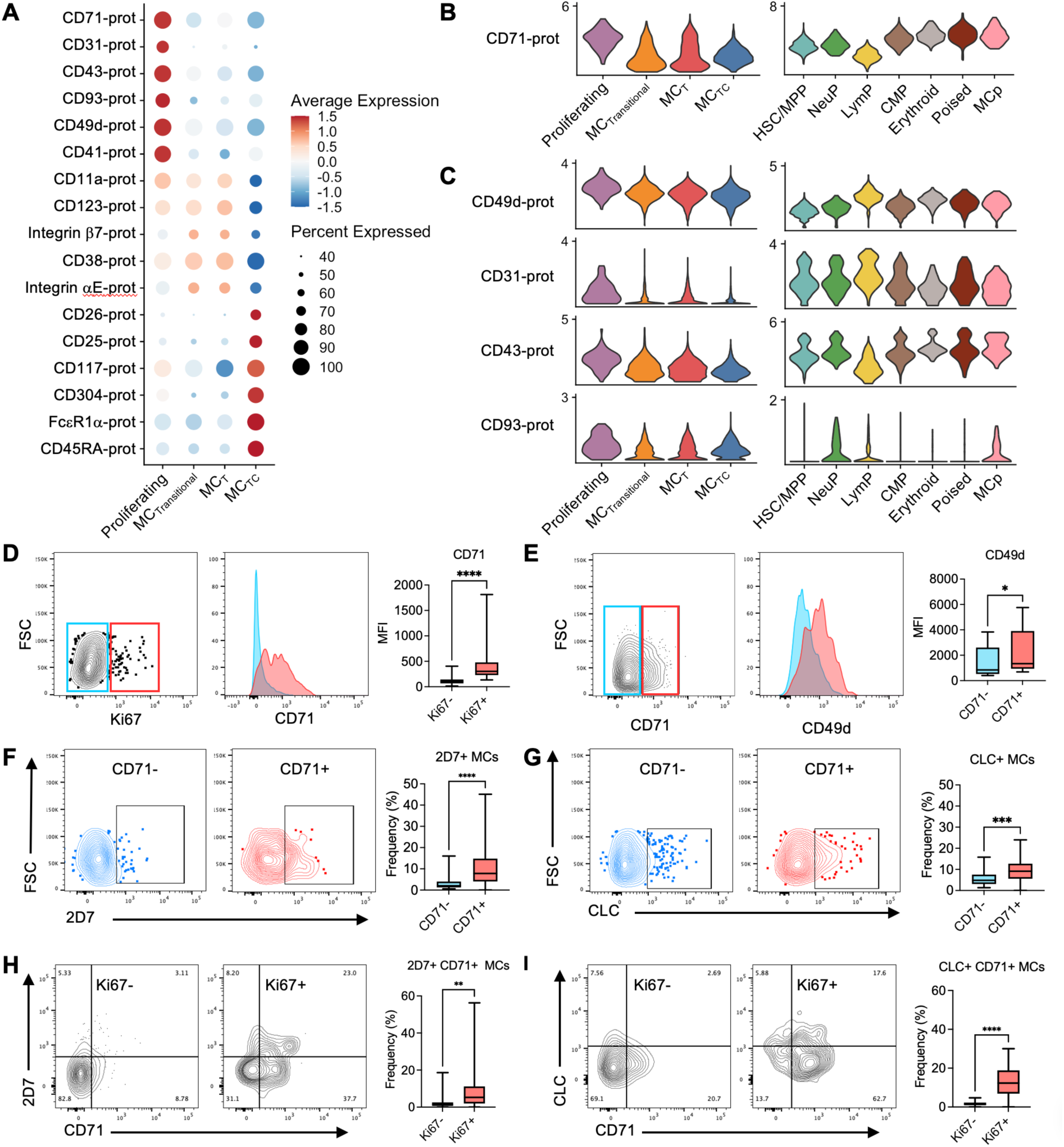
Proliferating tissue MCs exhibit a progenitor-like phenotype. A) Dot plot showing differentially detected cell surface proteins identified via CITE-seq across the tissue MC clusters. B-C) Violin plots showing select cell surface proteins significantly enriched in proliferating MCs via CITE-seq that were either significantly elevated (B) or highly expressed (C) in circulating MCp. D) Representative flow plots showing gating for Ki67^+^ (red) vs Ki67^-^ (blue) MCs (left), representative histogram showing CD71 expression (middle), and quantification of CD71 expression on Ki67^+^ vs Ki67^-^ MCs (right) (n=21 donors). E) Representative flow plots showing gating for CD71^+^ vs CD71^-^ MCs and histograms indicating CD49d expression with quantification (n=4 donors). F-G) Intracellular 2D7 (F) and CLC (G) content in CD71^+^ (red) and CD71^-^ (blue) MCs from CRSwNP donors (n=21 donors). H-I) Frequency of cells co-expressing 2D7 and CD71 (H) or CLC and CD71 (I) among proliferating (Ki67^+^, right) MCs relative to non-proliferating MCs (Ki67^-^, left) in CRSwNP tissue (n=21 donors). *, p < 0.05; **, p < 0.01; ***, p < 0.001; ****, p < 0.0001 (ratio-paired t test). For dot plots, Padj < 0.05 and Log2FoldChange > 0.5 for each subset relative to other tissue MC populations, color denotes cluster-averaged relative expression, size indicates percent of cells with marker detected.

To validate these observations in a larger sample cohort, we conducted a flow cytometric assessment of CRSwNP MCs. We observed a highly significant increase in cell surface CD71 expression on MCs containing nuclear Ki67, a marker of proliferation, relative to Ki67^-^ MCs (Fig. 3D). Further strengthening the link between circulating MCps and proliferating polyp MCs, CD71^+^ MCs exhibited significantly elevated cell-surface expression of CD49d (Fig. 3E), a marker previously associated with circulating MCps^5^, and were significantly enriched for intracellular 2D7 and CLC relative to CD71^-^ MCs (Fig. 3F-G). As in the circulation, basophils within the polyp (FcεR1α^+^ CD117^-^ CCR3^+^ 2D7^+^) contained bimodal intracellular levels of CLC but lacked CD71 expression, confirming that detection of CLC and 2D7 within the nasal polyp MC pool was not the result of basophil contamination (Fig. S4A). Previously, we observed that proliferating MCs in CRSwNP were enriched within the CD117^high^ CD38^high^ transitional MC gate and hypothesized that transitional MCs represented an immature population arising from recently recruited MCps^5^. Consistent with this hypothesis, MCs co-expressing CD71 with either 2D7 or CLC were significantly enriched within both the Ki67^+^ MC population and the transitional MC pool (Fig. 3H-I, S4B). MCs with this progenitor-like cell surface/intracellular phenotype were not present in sinonasal tissue from CRS patients lacking nasal polyps (CRSsNP) despite the presence of mature MCs, and no significant differences in MCp-like MCs were observed between CRSwNP and AERD (Fig. S4C). Overall, these data indicated that proliferating MCs in the nasal polyps were phenotypically linked with circulating MCps and enriched within the transitional MC pool.

This association between circulating MCps and proliferating polyp MCs was surprising based on their previously reported slow proliferation rate, with a reported one-week doubling time in ex vivo culture^6^. We previously observed that proliferating MCs in CRSwNP were primarily located within the subepithelial space^5^, potentially indicating that local tissue structural cells could regulate MCp proliferation. To test this hypothesis, MCps were flow-sorted from the peripheral blood of healthy donors and maintained for three weeks either in monoculture or co-culture with sinonasal fibroblasts from control donors, to avoid the potential confounding influence of disease heterogeneity (Fig. 4A). All cultures were supplemented with stem cell factor (SCF) to support viability. MCps in monoculture expanded ∼9-fold over the course of three weeks, consistent with their reported one-week doubling time. In contrast, MCps co-cultured with sinonasal fibroblasts exhibited a significant increase in proliferative capacity, instead expanding over 40-fold (Fig. 4B). CD45^+^ cells in monoculture and fibroblast co-cultures were uniformly positive for CD117 and FcεR1α and contained metachromatic granules, indicating pure MC populations (Fig. 4C-D). Fibroblasts have previously been found to influence MC expansion and maturation through trans-presentation of membrane-bound SCF^18,19^. However, no reduction in MCp proliferation was observed when MCps were separated from fibroblasts by a transwell insert, indicating that a soluble factor was responsible for the enhanced MCp proliferation (Fig. 4E). Thus, MCps rapidly proliferate and mature following exposure to sinonasal fibroblasts, potentially explaining why so few events transcriptionally resembling circulating MCps were observed in CRSwNP tissue (Fig. 1F).

**Figure 4:**
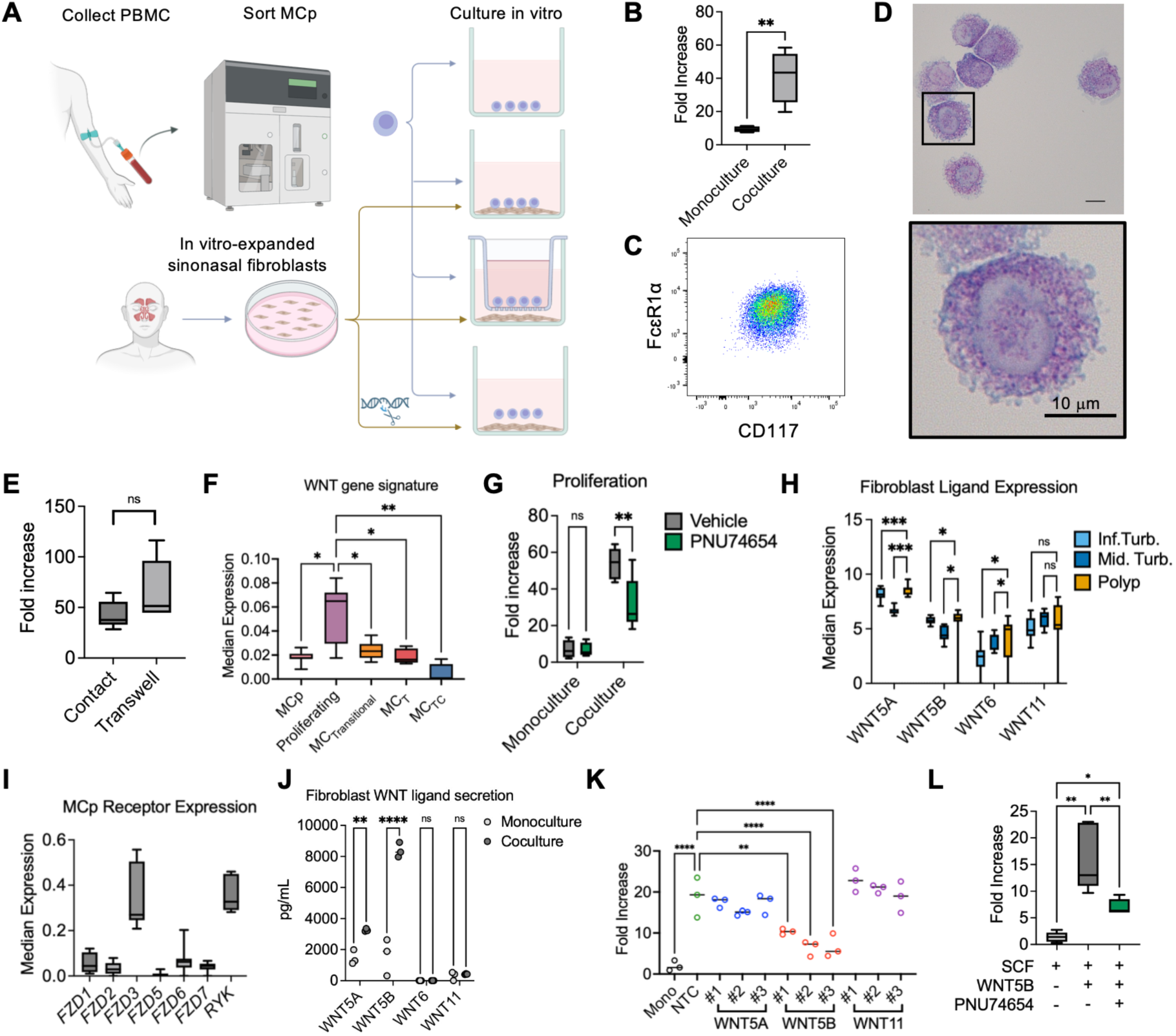
Fibroblasts direct WNT5B-dependent MCp proliferation. A) Schematic representation of ex vivo MCp culture system. B) Quantification of fold-change expansion of 200 MCp in monoculture or co-culture with sinonasal fibroblasts for three weeks supplemented with 20 ng/ml SCF (n=5 donors). C) Representative flow showing CD45^+^ cell expression of FcεR1α and CD117 following three weeks co-culture with sinonasal fibroblasts. D) Toluidine blue staining of flow-sorted CD45^+^ cells following fibroblast co-culture, with inset magnified. Scale bar indicates 10 μm. E) Quantification of MCp proliferation in direct co-culture with fibroblasts or following separation by a transwell membrane (n=5 donors). F) Donor-averaged quantification of WNT signature expression as a percentage of all transcripts across circulating MCp and sinonasal MC subsets. G) Quantification of MCp expansion in monoculture or co-culture with fibroblasts following treatment with vehicle or the β-catenin inhibitor PNU74654 (n=4-5 donors/condition). H) Donor-averaged WNT ligand expression on sinonasal fibroblasts from polyp tissue versus fibroblasts from inferior and middle turbinate tissues. I) Donor-averaged expression of WNT receptor transcripts in circulating MCps. J) Quantification of soluble WNT ligand detection in supernatants of sinonasal fibroblasts maintained in monoculture or in co-culture with circulating MCps for one week (n=3 donors). K) Quantification of MCp expansion in co-culture with sinonasal fibroblasts following CRISPR/Cas9-mediated deletion of the indicated WNT ligands using three independent guide RNAs for each ligand (n=3 donors). L) MCp expansion in monoculture following treatment with SCF, SCF plus WNT5B, or SCF and WNT5B plus PNU74654 (n=5 donors). *, p < 0.05; **, p < 0.01; ***, p < 0.001; ****, p < 0.0001 (ratio-paired t test for pairwise comparisons, ANOVA for multi-group comparisons).

In contrast to prior observations that MCp concentration was elevated in the circulation of asthmatics^6^, we did not observe any differences in circulating MCp concentration between healthy control donors, CRSsNP donors, aspirin-tolerant CRSwNP donors, or patients with AERD, either as a proportion of PBMCs or as a fraction of CD34^+^ cells (Fig. S5A). Similarly, no MCp-intrinsic differences in proliferative capacity were observed across groups (Fig. S5B). Thus, we hypothesized that the increased MCp proliferation observed in CRSwNP likely arose from differences in MCp recruitment to the tissue and/or local fibroblast-derived signals rather than MCp-intrinsic differences across disease groups.

Previously, we found that transitional MCs in AERD nasal polyps were enriched for a gene signature of Wingless/Integrated (WNT) signaling^10^, and murine gain-of-function studies have linked WNT signaling with tissue MC hyperproliferation^20^. We observed significant enrichment for a cross-tissue signature of human WNT target genes^21^ in proliferating MCs in our scRNA-seq dataset relative to both MCps and mature tissue MC populations (Fig. 4F). As airway fibroblast-derived WNT ligands have previously been implicated in modulating epithelial cell function^22^, we assessed whether fibroblast amplification of MCp proliferation may similarly be WNT-dependent by inhibiting β-catenin, a canonical mediator of WNT signaling. Treatment with the β-catenin inhibitor PNU74654 had no impact on MCp proliferation in monoculture but significantly reduced MCp proliferation in co-culture with fibroblasts, albeit without completely negating the impact of fibroblasts (Fig. 4G). Thus, both in vitro co-culture data and in vivo transcriptional signatures identified fibroblast-driven WNT signaling as a potential driver of MCp proliferation.

We next assessed in vivo sinonasal fibroblast expression of WNT ligands using a recently published scRNA-seq study of nasal polyp versus turbinate tissue in CRSwNP patients (Fig. S6A-B)^23^. Among ligand-encoding transcripts detected, *WNT5A*, *WNT5B*, *WNT6* and *WNT11* stood out as robustly expressed in polyp fibroblasts, with *WNT5A*, *WNT5B* and *WNT6* exhibiting significant increases within the fibroblast cluster in polyp tissue relative to turbinate tissue (Fig. 4H). A sub-analysis of fibroblasts indicated substantial tissue-associated differences in the fibroblast compartment, with three polyp-enriched clusters, four turbinate-enriched clusters, and two mixed clusters (Fig. S6C-E). Among polyp-enriched fibroblasts (clusters P1-P3), the P1 and P2 clusters were notably marked by robust expression of *OXTR*, encoding the oxytocin receptor, a marker previously identified on a population of tumor-associated fibroblasts linked with aggressive tumor invasion and metastasis^24^, while *POSTN*, encoding periostin, was observed in all polyp fibroblast clusters (Fig. S6E). *WNT5A* was significantly enriched across all polyp-associated fibroblast clusters, while *WNT5B* was enriched in the P1 and P2 clusters and *WNT6* was specifically enriched in the P1 cluster (Fig. S6F). Analysis of circulating MCps revealed detection of receptors associated with each ligand (WNT5A: FZD1, FZD2, FZD3, FZD5, FZD6, FZD7, Ryk; WNT5B: FZD1, FZD6, Ryk; WNT6: FZD2; WNT11: FZD7)^25,26^ (Fig. 4I). As WNT ligands can be either membrane-bound or soluble, we evaluated the concentration of each ligand in the supernatant of ex vivo-cultured sinonasal fibroblasts, in monoculture or co-culture with MCps. WNT5A, WNT5B and WNT11 were all detected in the supernatant, while WNT6 was undetectable (Fig. 4J). Notably, both WNT5A and WNT5B levels were significantly increased following MCp co-culture, with WNT5B exhibiting the most robust induction.

To evaluate the role of each candidate ligand in driving MCp proliferation, we conducted CRISPR/Cas9-mediated gene deletion using three guide RNAs directed against each of the three candidates detected in culture supernatants (WNT5A, WNT5B, WNT11), confirming significant reductions in all targets via immunoblotting (Fig. S7). Fibroblast-dependent MCp expansion was significantly reduced following co-culture with WNT5B-edited fibroblasts for all guide RNAs relative to co-culture with non-targeting control (NTC)-treated fibroblasts, while no impact on MCp proliferation was observed following co-culture with fibroblasts in which WNT5A or WNT11 were targeted (Fig. 4K). As WNT factors can exhibit both autocrine and paracrine effects, and autocrine roles for WNT5A in fibroblasts have previously been reported^27,28^, we evaluated whether WNT5B alone was sufficient to drive MCp proliferation. Indeed, MCps cultured with SCF plus WNT5B exhibited a significant increase in proliferation relative to MCps cultured in SCF alone (Fig. 4L). Although WNT5B is reported to primarily signal through a non-canonical WNT signaling pathway^26^, β-catenin inhibition significantly blunted the amplifying effect of WNT5B on MCp proliferation (Fig. 4L), indicating that WNT5B likely engages both canonical and non-canonical signaling pathways in MCps. Interestingly, a prior murine study similarly found that the non-canonical WNT ligand WNT5A elicited murine bone marrow-derived MC maturation through the canonical β-catenin-dependent pathway^29^, suggesting a broader paradigm of non-canonical WNT ligands engaging the canonical pathway in MCs. Within polyp tissue, *WNT5B* expression was also detected within the epithelial, myeloid and plasma cell clusters; thus, we cannot exclude the possibility that other cellular sources of WNT5B may also contribute to MC proliferation in CRSwNP (Fig. S6G).

Together, our findings link circulating MCps with proliferating MCs in CRSwNP polyp tissue and demonstrate that MCps exhibit robust proliferative potential, helping to explain the disconnect between the scarcity of circulating MCps and the relative abundance of mature MCs under inflammatory conditions. We determined that tissue microenvironmental cues beyond SCF can regulate MC expansion, identifying fibroblast-derived WNT5B as a novel driver of MCp proliferation. In recent years, fibroblasts have emerged as key regulators of tissue immunity during homeostasis and inflammation^30^. In the lower airways, fibroblasts were recently found to localize to MC/macrophage inflammatory hubs in asthma^31^. Combined with our findings highlighting MCp/fibroblast crosstalk as a key regulatory circuit directing MC hyperplasia during CRSwNP, these observations support a broader paradigm of MC/fibroblast interactions playing a key role in human airway disease.

## Methods

### Sex as a biological variable

The study was conducted using samples from both female and male donors based on sample availability. Sex was not assessed as a biological variable.

### Ethical statement for human experiments

All human experiments were approved by the Brigham and Women’s Hospital Institutional Review Board (protocol no. 2013P001659), and all study participants provided written consent at the Brigham and Women’s Hospital. All study procedures were conducted in accordance with the guidelines of the institutional ethics committee. Ethmoid sinus tissue and peripheral blood were collected at the time of elective endoscopic sinus surgery from patients with physician-diagnosed CRSwNP or CRSsNP. CRSwNP patients included a mix of aspirin-tolerant CRSwNP and aspirin-intolerant AERD. A diagnosis of AERD was suspected for patients with asthma, nasal polyposis, and a history of respiratory reaction following use of cyclooxygenase-1 inhibitors, and AERD diagnoses were confirmed via graded oral challenge with aspirin. All tissues were obtained from participants between the ages of 20 and 80 years recruited from the Brigham and Women’s Hospital Allergy and Immunology clinics and Otolaryngology clinics.

### Antibodies

The following anti-human antibodies were purchased from BioLegend. For flow cytometry: CD117 (104D2), FcεR1α (CRA-1), CD45 (2D1), CD11b (ICRF44), CD11c (3.9), CD38 (HB-7), CD3 (OKT3), CD19 (HIB193), CD14 (M5E2), CD56 (5.1H11), CD84 (CD84.1.21), CD33 (WM53), CD45RA (HI100), GPR56 (4C3), HLA-DR-DP-DQ (TU39), IL-3Rα (6H6), CD116 (4H1), CD203c (NP4D6), CD131 (1C1), CD71 (CY1G4), CCR3 (5E8), 2D7 (Basophil/2D7), CD49d (9F10), Ki67 (Ki-67), CD90 (5E10), and mouse IgG1 isotype control (MOPC-21). MS4A3 (REA756) and CD34 (AC136) were purchased from Miltenyi Biotec. CLC/Galectin-10 (B-F42) was purchased from Diaclone and labeled using an Alexa Fluor 647 antibody labeling kit (Invitrogen).

For CITE-seq, the following TotalSeqB clones were purchased and used in all experiments: CD117 (104D2), Integrin αE (Ber-ACT8), Integrin β7 (FIB504), FcεR1α (CRA-1), CD38 (HB-7), CD49d (9F10), CD133 (S16016B), CXCR4 (12G5), and CD71 (CY1G4). The TotalSeqB Human Cocktail v1.0 was additionally used for four donors. For experiments in which the cocktail was not used, biotin-conjugated anti-CD131 (1C1) was used in conjunction with TotalSeqB0975 Streptavidin.

### Flow cytometry

PBMCs or dissociated ethmoid sinus cells were incubated with human TruStain FcX (BioLegend) prior to staining to block nonspecific antibody binding. Cell surface markers were stained for 40 min on ice. For detection of intracellular markers (CLC, 2D7, Ki67), cells were fixed and permeabilized using the BioLegend TrueNuclear transcription factor staining kit following the manufacturer’s protocol. Cells were incubated in blocking buffer containing 1% BSA and human TruStain FcX for at least 15 min prior to staining for intracellular markers for 30 min at room temperature. Anti-human CLC was conjugated to AF647 following the manufacturer’s protocol (Invitrogen). Cell viability was measured by staining with a viability staining solution (Zombie fixable viability dye or 7-AAD, BioLegend). For conventional flow cytometry, cells were collected on a BD FACSCanto II or BD LSRFortessa using BD FACSDiva software. For flow cytometry-based sorting, samples were collected using a Sony SH800S cell sorter.

### Primary human sample processing

For peripheral blood mononuclear cell (PBMC) isolation, peripheral blood was collected into 10 ml Vacutainers (Becton Dickinson) containing sodium heparin. Whole blood was diluted 1:1 with HBSS -/-containing 2 mM EDTA, layered on top of Ficoll-Paque Plus (GE Healthcare), and centrifuged for 30 min at 500 RCF without brake. The interface was collected and washed twice with HBSS -/-containing 2 mM EDTA at 110-280 RCF to remove platelets, after which any remaining erythrocytes were lysed using ACK buffer.

Surgical specimens were collected into 30 mL of ice-cold RPMI (Corning) and digested as previously described^32^. Briefly, specimens were finely minced between two scalpel blades and incubated for 15 min at 37°C in a rotisserie rack with end-over-end rotation in 25 ml digestion buffer supplemented with 600 U/ml collagenase IV (Worthington) and 20 µg/ml DNase I (Roche) in RPMI with 10% fetal calf serum (FCS). After 15 min, samples were triturated five times using a syringe with a 16G needle and returned to the rotisserie rack for another 15 min. After the second digest period, samples were triturated an additional five times using a syringe with a 16G needle, at which point the digestion was stopped via the addition of EDTA to a final concentration of 20 mM. Samples were typically fully dissociated at this step and were filtered through a 70 µm cell strainer and spun down at 500 RCF for 10 min followed by a rinse with ice-cold PBS (Thermo Fisher, Ca/Mg free) in 30 ml total volume. To maintain consistency across patient groups, erythrocytes were lysed using ACK buffer (Thermo Fisher) for 3 min on ice, even if no contamination was visibly seen.

### Primary human cell culture

MCps were flow-sorted from PBMCs of healthy human volunteers using a Sony SH800S cell sorter based on a Lineage (CCR3, CD3, CD11b, CD11c, CD14, CD19, and CD56)^-^ CD34^+^ CD131^+^ CD71^+^ CD117^+^ FcεR1α^+^ cell surface profile. Primary human sinonasal fibroblasts (passage 2 to 4) were seeded at 10,000 cells/well along with isolated MCps at 200 cells/well in StemSpan SFEM (STEMCELL Technologies) supplemented with 100 U/ml penicillin, 100 µg/ml streptomycin, and 20 ng/ml rhSCF (Peprotech) to support viability. Each week, half of the culture media was removed and replaced. After three weeks, cultures were harvested and assessed via flow cytometry, with dead cells excluded from analysis using 7-AAD or DAPI. WNT/β-catenin signaling was assessed using the β-catenin inhibitor PNU74654 (Cayman Chemical) or vehicle control added at 125 µM. In co-culture experiments assessing WNT ligands, StemSpan media was supplemented with 100 U/ml penicillin, 100 µg/ml streptomycin, 200 nM 2-mercaptoethanol and 10 ng/ml rhSCF. Sinonasal fibroblasts were seeded at 40,000 cells/well two days prior to the addition of freshly isolated MCps, with media changed weekly. In some experiments, MCp monocultures were treated with 100 ng/ml rhWNT5B (R&D Systems) or vehicle control.

### ELISA

The concentrations of WNT5A, WNT5B, WNT6, and WNT11 were assessed in the supernatant of primary human sinonasal fibroblast cultures via ELISA using kits purchased from antibodies-online according to the manufacturer’s instructions. Fibroblasts were maintained for one week in monoculture or co-culture with MCps flow-sorted from the peripheral blood of healthy human volunteers, after which supernatants were collected for ELISA analysis.

### Immunoblotting

For immunoblotting detection of WNT5A, WNT5B, and WNT11 in the whole cell lysate of the primary human sinonasal fibroblasts, 5 x 10^5^ cells were lysed with 200 µL of CelLytic M (Sigma-Aldrich) treated with protease inhibitor cocktail cOmplete ULTRA Tablets (Roche), and denatured at 65°C for 7 min (WNT5A), 70°C for 3 min (WNT5B and WNT11), or 95°C for 5 min (β-actin). After SDS-polyacrylamide gel electrophoresis (SDS-PAGE), separated proteins were transferred onto the polyvinylidene difluoride (PVDF) membranes by electroblotting, incubated with 5% bovine serum albumin (BSA), and immunoblotted with rabbit anti-human WNT5A monoclonal antibody (23GB2400), rabbit anti-human WNT5B monoclonal antibody (JE53-67), or rabbit anti-human WNT11 monoclonal antibody (HL1630) all from Invitrogen, or Horseradish peroxidase (HRP)-conjugated rabbit anti-human β-actin monoclonal antibody (13E5, Cell Signaling Technology). WNT ligands and β-actin loading controls were assessed on parallel membranes due to differences in the denaturing conditions required for each target. For the detection of each WNT, HRP-conjugated donkey anti-rabbit IgG polyclonal antibody (BioLegend) was then incubated for 30 min with gentle shaking, and HRP was detected by ECL western blotting substrate (Thermo Fisher Scientific) using a ChemiDoc MP Imaging System (Bio-Rad).

### Single-cell RNA sequencing

Peripheral blood and sinus tissues were obtained from Brigham and Women’s Hospital. Single-cell suspensions were stained for flow cytometry analysis. TotalSeq hashtag antibodies (BioLegend) were added for sample multiplexing and for tracing sample origins, together with TotalSeqB CITE-seq antibodies directed against CD117, Integrin αE, Integrin β7, FcεR1α, CD38, CD49d, CD133, CD131, CXCR4, and CD71. For a subset of patients, the BioLegend TotalSeqB human cocktail v1.0 was added. Dead cells were excluded using propidium iodide or Zombie viability dye (BioLegend). Sorted cells were resuspended in 0.4% BSA in PBS at a concentration of 1,000 cells per μl, and loaded onto a single lane of a Chromium chip (10X Genomics) for encapsulation using the single cell 3′ kit V3.1 (10X Genomics). cDNA synthesis and gene expression and protein library generation were performed according to the 10X Genomics protocol. The generated gene expression libraries were sequenced to an average of 30,000 reads per cell and the surface protein libraries to an average of 15,000 reads per cell using the Illumina NovaSeq platform. scRNA-seq reads were aligned to the GRCh38 assembly using the STAR aligner implemented within Cell Ranger to quantify transcript counts per putative cell.

Prior to downstream analysis, count matrices were processed using CellBender to correct for background ambient RNA and oligo-tagged antibodies. The filtered matrices were individually imported into Seurat v5^33^. For each sample, a dynamic upper limit for detected genes and unique molecular identifiers was set at the 75th percentile + 1.5 × the interquartile range on a per-sample basis, and a lower limit of 350 genes and 600 unique molecular identifiers was used, with all cells not meeting these criteria filtered out. Cells with greater than 20% mitochondrial genes were excluded, after which matrices were merged prior to downstream analysis. Data from the eight donors were integrated using the Harmony package for R using default settings^34^. A shared nearest neighbor (SNN) graph and uniform manifold approximation and projection (UMAP) embedding were constructed using the first 50 Harmony components. The FindClusters function was used with the default resolution of 0.8 and RunUMAP used with the default settings to identify 20 clusters across the eight input samples. Clusters containing MC genes or *CD34* and consistently observed across all donors without other contaminating populations were computationally subsetted, after which data were re-integrated using Harmony with the same settings as before. The first 8 harmony components were chosen to construct a SNN graph and UMAP embeddings following visual inspection of the elbow plot, and clusters were determined at a resolution of 0.6 based on stability after evaluating all resolutions between 0.2 and 2 using the clustree package for R^35^. Quality control plots and violin plots were generated using the scCustomize package for R^36^.

Differential gene expression analysis between clusters was conducted using a pseudobulk approach, based on recent benchmarking studies^37^. Raw counts within each cluster were aggregated by donor, after which differential gene expression analysis was conducted using DESeq2, contrasting each cluster to the average of all other clusters. All transcripts with fewer than 2,500 total counts across all replicates were removed from analysis. Transcripts identified as differentially expressed between clusters with an adjusted p value of less than 0.05 were considered statistically significant.

For re-analysis of existing human sinonasal fibroblasts, count matrix files from Xie et al^23^ were obtained from the NIH Gene Expression Omnibus (Accession GSE276503) and analyzed using Seurat via the pipeline described above, with analysis restricted to donors with CRSwNP. Fibroblasts were selected based on canonical marker gene expression profile. After subsetting, the first 16 harmony components were chosen to construct a SNN graph and UMAP embeddings following visual inspection of the elbow plot, and clusters were determined at a resolution of 0.5 based on stability after evaluating all resolutions between 0.1 and 1 using the clustree package for R^35^.

### CRISPR-Cas9 editing of WNT ligands in human sinonasal fibroblasts

Primary sinonasal fibroblasts were cultured in DMEM culture medium supplemented with 10% heat-inactivated FCS, 2 mM glutamate, 100 U/ml penicillin, 100 µg/ml streptomycin, and 200 nM 2-mercaptoethanol. Thirty pM of single guide (sg) RNAs directed against WNT5A, WNT5B, and WNT11, using three separate sgRNA per target, were incubated with GFP-tagged Cas9 nuclease (#10008100, Integrated DNA Technologies) for 10 min at room temperature with light protection, followed by transfection of the sgRNA-Cas9 complex through electroporation. After transfection, cells were resuspended in D-MEM culture medium supplemented with 10% heat-inactivated FCS, 3 mM glutamate, 1 µM HEPES, 20 µM sodium pyruvate, 100 µM MEM non-essential amino acids, and 200 nM 2-mercaptoethanol, without penicillin or streptomycin, for 10 min under 5% CO_2_ at 37°C, after which culture medium was replaced with D-MEM culture medium supplemented with 15% heat-inactivated FCS, 3 mM glutamate, 100 U penicillin, 100 µg/ml streptomycin, 1 µM HEPES, 20 µM sodium pyruvate, 100 µM MEM non-essential amino acids, and 200 nM 2-mercaptoethanol. Sequences for sgRNAs were as follows: WNT5A: #1 CGAGACGGCCTTCACATACG, #2 AGTATCAATTCCGACATCGA, #3 GAGGATGCGAGCACTCTCGT; WNT5B: #1 CGGACAACGCATCTGTCTTT, #2 CTGCAAATGCCACGGCGTCT, #3 GCACCGATGATAAACATCTC; WNT11: #1 CATACACGAAGGCCGACTCC, #2 TGTGCGGACAACCTCAGCTA, #3 GTCCAAGACACCATCGGCCC. According to the manufacturer’s instructions, the non-targeting control crRNA was annealed with tracrRNA at 95°C for 5 min before transfection.

### Statistics

For experiments not involving transcriptomics, data are presented as mean ± SEM or with box-and-whisker plots displaying median, interquartile range and minimum/maximum values. Statistical analyses were conducted using the ratio-paired t test for paired samples or ANOVA for analyzing three or more groups using GraphPad Prism 10.6.1 (GraphPad Software), with p values less than 0.05 after correcting for multiple comparisons considered statistically significant. For analysis of scRNA-seq and CITE-seq, adjusted p values of less than 0.05 were considered as significant, with additional cutoff values of absolute Log_2_FoldChange > 0.5 where indicated.

## Acknowledgments

This work was supported by NIH grants R21 AI180332, 2, R01 AI078908, R37 AI052353, R01 AI136041, R01 AI130109, R01 AI175149, U19 AI095219, and by generous contributions from the Vinik family. We thank the Brigham and Women’s Hospital Center for Cellular Profiling for RNA-seq support, and Daniel Wing for administrative support.

## Author contributions

T.D., J.B., and D.D. conceived the study. T.D., K.M., and D.D. designed experiments, analyzed data and interpreted results. T.D., K.M., R.S., E.H., C.N., and A.P. conducted experiments. R.K. assisted with transcriptomics analysis. S.K. assisted with implementation and troubleshooting for CRISPR/Cas9 editing. K.B. and T.L. oversaw and coordinated human sample collection. Sinus surgeries were conducted by R.B., S.L., A.M., and R.R. T.D., K.M. and D.D. wrote the manuscript, with input from all authors.

## Data and materials availability

Raw count matrices and processed Seurat object for the single-cell RNA-Seq datasets of circulating progenitors and mature polyp MCs will be made available prior to final publication.

