## Supplementary material for "Airway fibroblasts drive WNT5B-dependent mast cell progenitor proliferation": Data supplement

**Supplemental Figures**

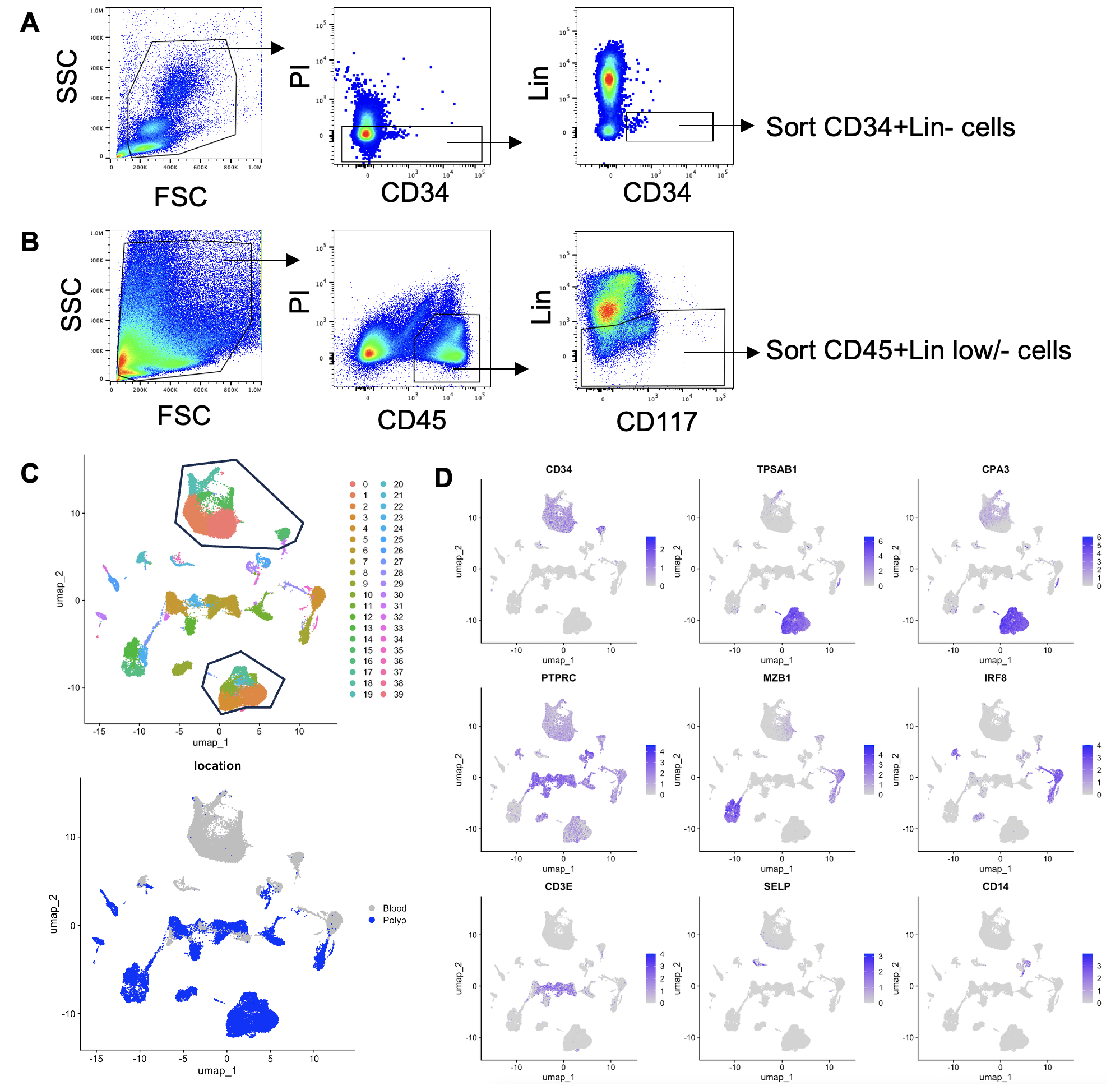
**Figure S1: Generation of a multimodal single-cell dataset of nasal polyp and peripheral blood mononuclear cells.** A) Flow scheme for isolating CD34^+^Lin^-^ cells from peripheral blood. B) Flow scheme for isolating CD45^+^Lin^-^ cells from CRSwNP digested tissue. C) Uniform Manifold Approximation and Projection (UMAP) embedding for cells passing quality control from eight donors (top) and location-of-origin for each cell (bottom). D) Marker gene expression for circulating progenitors, MC lineage cells, and other populations observed.

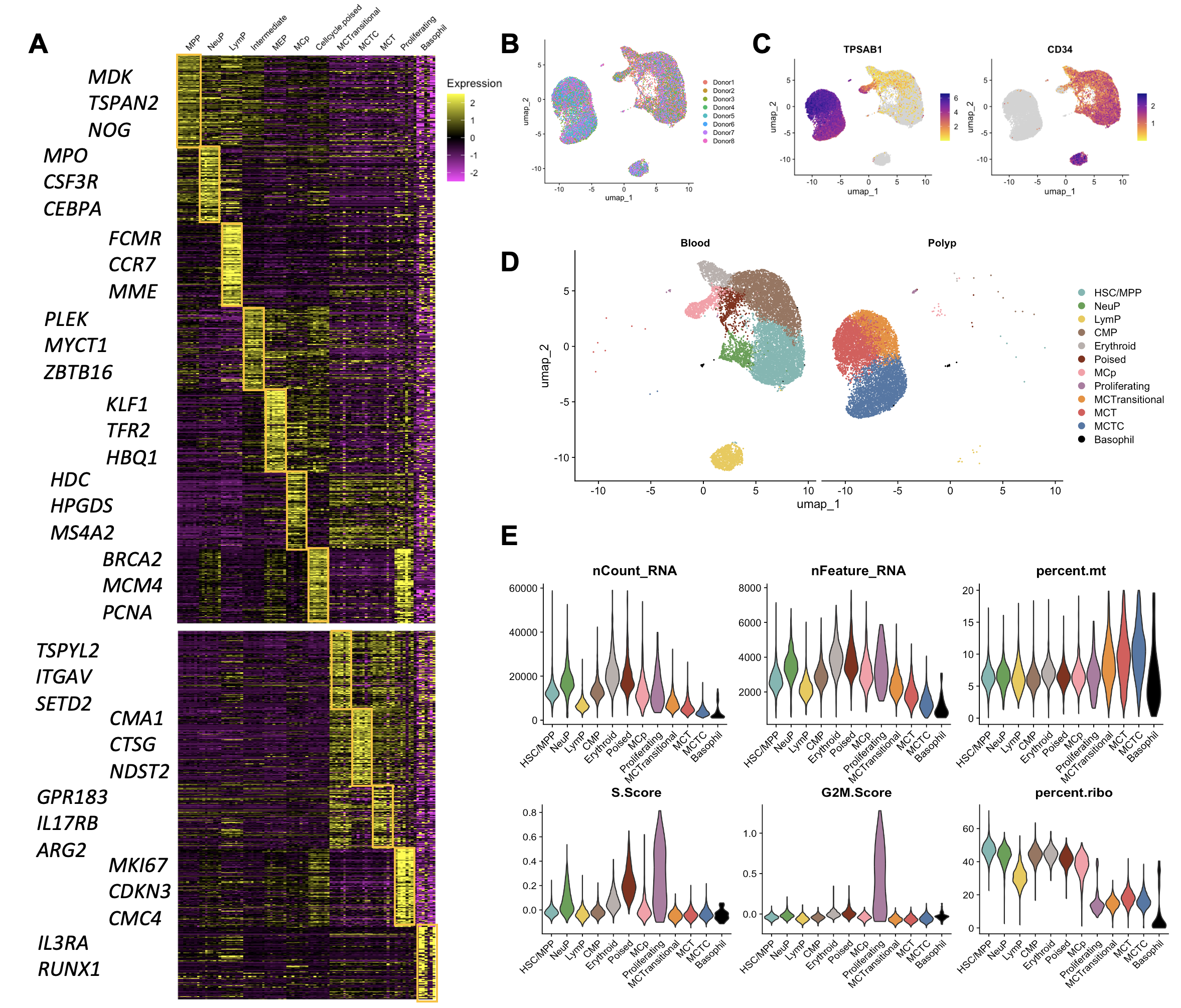
**Figure S2: Characterization of circulating progenitors and tissue-associated MCs.** A) Donor-averaged heatmap showing top 100 differentially expressed transcripts across clusters with select genes highlighted. B) UMAP display colored by individual donor of origin. C) Transcript expression for *TPSAB1* and *CD34* across all cells. D) UMAP display of identified clusters stratified based on hashtag expression. E) Violin plots showing QC metrics and cell cycle analysis across all clusters.

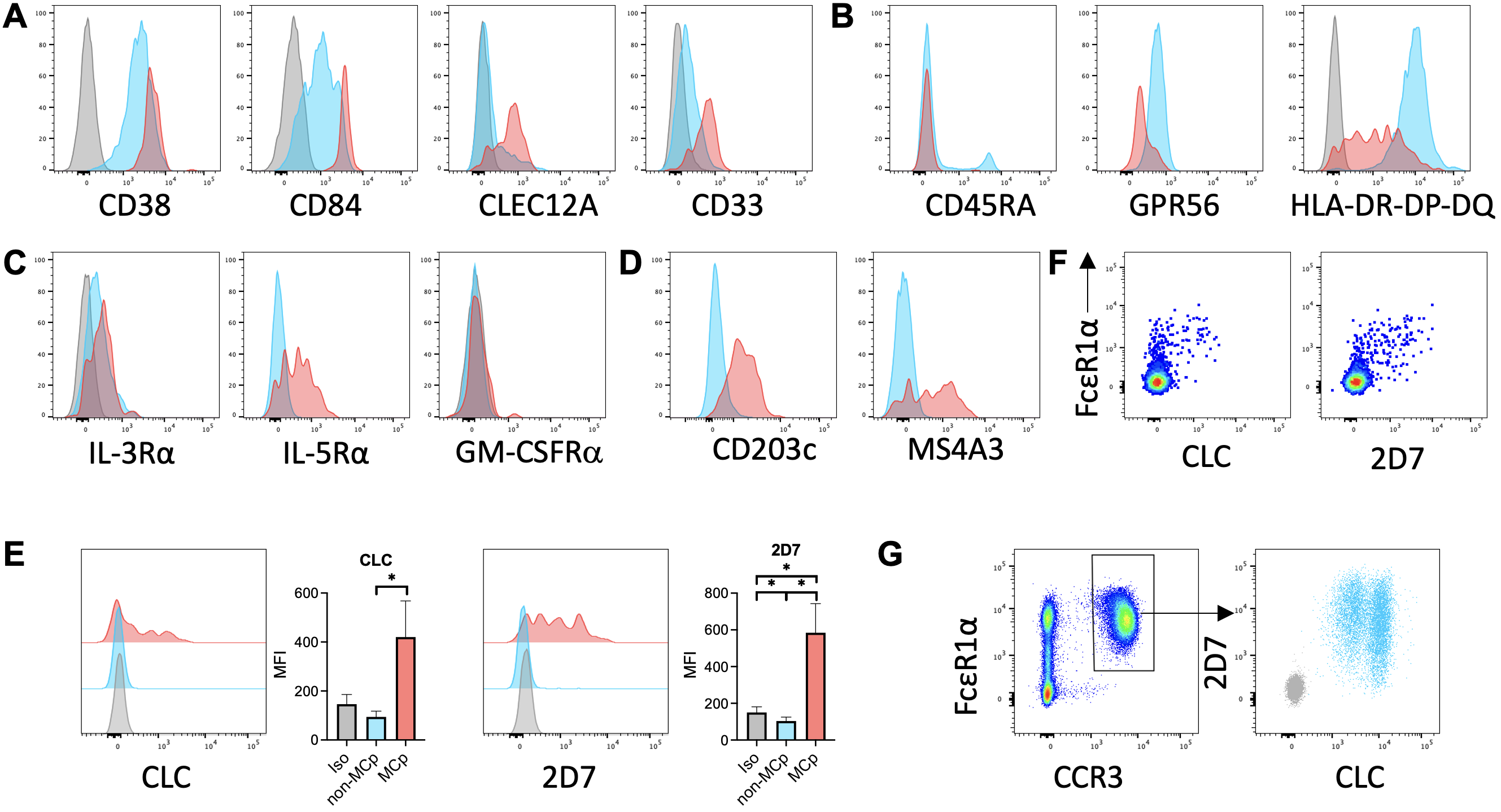
**Figure S3: Flow cytometric evaluation of MCp cell surface protein expression.** A-B) Flow validation of select proteins identified through CITE-seq as (A) enriched on MCps or (B) exhibiting diminished expression on MCps (red) relative to other circulating progenitors (blue) and isotype control (grey). C) Histograms showing expression of CD131-paired receptors on MCps (red) vs other circulating progenitors (blue) and isotype (grey). D) Histograms showing expression of CD203c and MS4A3, encoded by the MCp-enriched transcripts *ENPP3* and *MS4A3*, respectively. E) Quantification of CLC (left) and 2D7 (right) expression in MCp relative to all other circulating CD34^+^ cells. F) FcεR1α expression relative to intracellular CLC and 2D7 in all CD34^+^ cells. G) 2D7 vs CLC expression in circulating basophils.

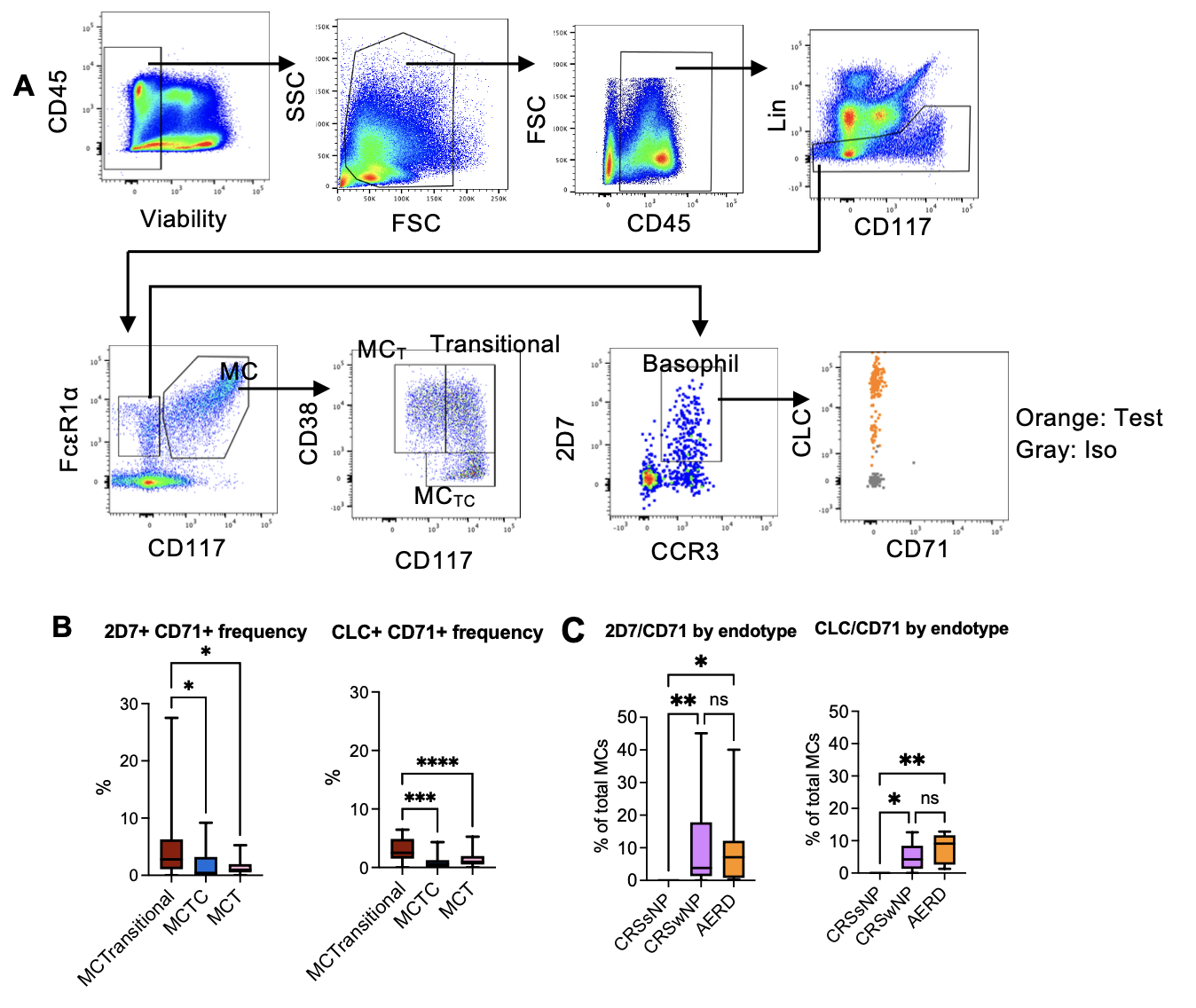
**Figure S4: MCp and basophil comparison in CRSwNP tissue.** A) Gating scheme for identifying MC and basophil subsets in CRSwNP tissue**.** Arrows indicate sequential gates. For basophil CD71 staining, orange indicates stain, grey indicates isotype. B) Quantification of MCs co-expressing CD71 and either 2D7 or CLC among flow-defined polyp MC subsets identified as shown in (A). C) Quantification of MCs co-expressing CD71 and either 2D7 or CLC across chronic rhinosinusitis endotypes.

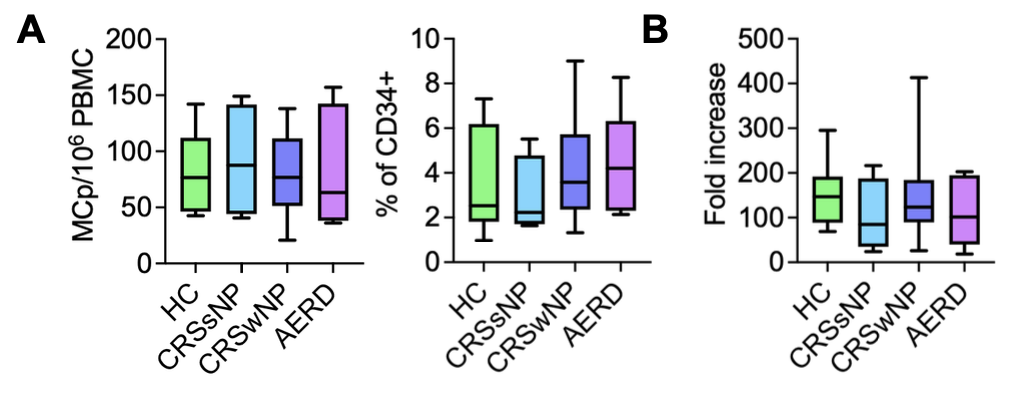

**Figure S5: MCp concentration and proliferation capacity across patient populations.** A) Quantification of MCps as percent of CD34^+^ cells or relative to all PBMCs across the indicated patient endotypes (n=4-15 donors per group). B) Quantification of MCp expansion potential across patient endotypes after co-culture with sinonasal fibroblasts (n=4-15 donors per group).

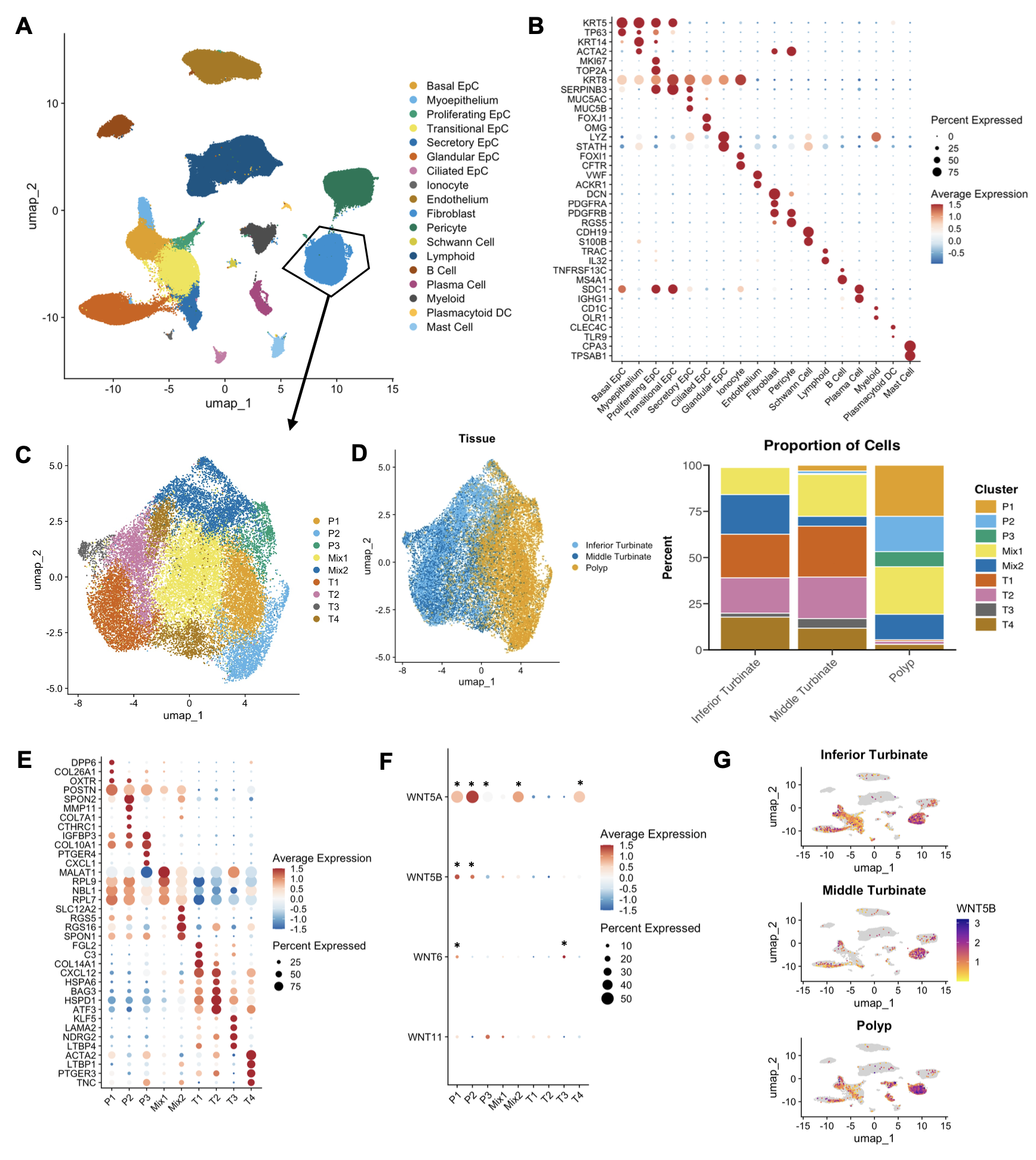

**Figure S6: scRNA-seq identification of fibroblasts in CRSwNP.** A-B) The sinonasal dataset generated by Xie et al^23^ was re-analyzed after subsetting on CRSwNP donors, and stromal and immune populations identified through canonical marker gene expression. C) Subset analysis of fibroblasts, identifying nine clusters. D) Quantification of clusters in turbinate versus polyp tissue (left) and frequency distribution of cells within each cluster across the three tissue compartments sampled (right). E) Dot plot showing enriched transcripts associated with each of the nine fibroblast clusters. F) Quantification of WNT ligand expression across the nine fibroblast clusters, * indicates significant enrichment (Padj < 0.05). G) WNT5B expression patterns in inferior (top) and middle (center) turbinate vs polyp (bottom).

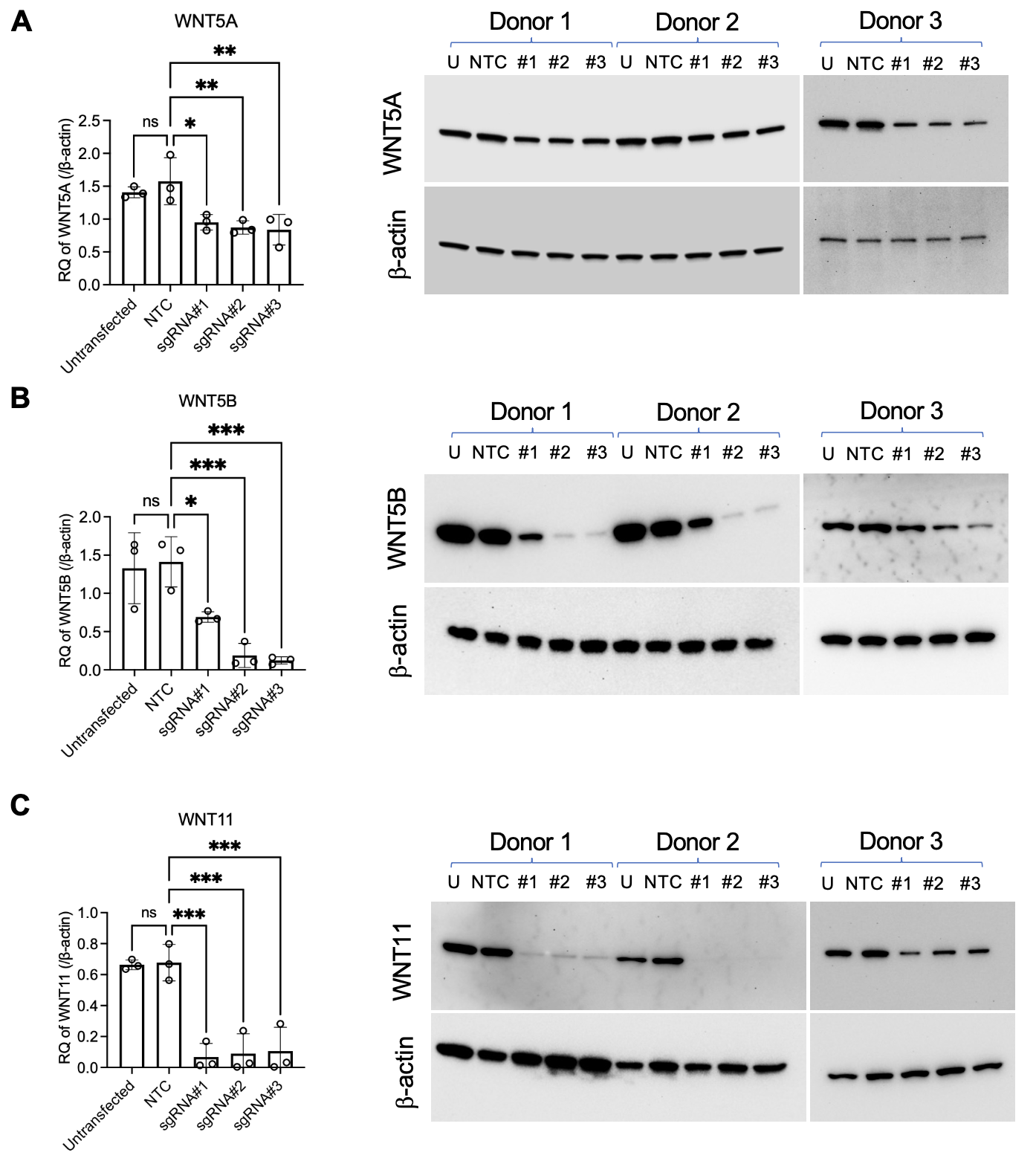

**Figure S7: Fibroblast regulation of MC proliferation.** Quantification and immunoblots showing protein content of A) WNT5A, B) WNT5B and C) WNT11 (top row) and β-actin loading controls (bottom row) in unedited fibroblasts (U) or following CRISPR/Cas9 editing using non-targeting control guide RNA (NTC) or three target-specific guide RNAs (#1, #2, #3). WNT ligand blotting and β-actin blotting were immunoblotted on separate gels run in parallel due to different denaturing conditions required for each target (WNT5A: 65°C for 7 min; WNT5B and WNT11: 70°C for 3 min; β-actin: 95°C for 5 min). *, p < 0.05; **, p < 0.01; ***, p < 0.001 (ANOVA).
